# Gut microbiome variation in juvenile blue tits in a European urban mosaic

**DOI:** 10.1101/2025.03.26.643606

**Authors:** Lena Fus, Sebastian Jünemann, Irene Di Lecce, Joanna Sudyka, Marta Szulkin, Öncü Maraci

## Abstract

Urbanisation transforms natural environments, impacting not only wild animals living in cities but also the microorganisms they are hosting. To better understand urban-driven variation in microbiological composition and diversity in the gut of birds developing in urban areas, we collected faecal samples from blue tit *Cyanistes caeruleus* nestlings using nestboxes distributed across the capital city of Warsaw, Poland. Sampling included a variety of urban habitats, a suburban village and a natural forest area. Microbiome analysis unveiled a pattern of reduced alpha and beta diversity in urbanised settings driven by impervious surfaces. Additionally, we observed that this effect was year-dependent, therefore highlighting the importance of temporal replication in ecological research. Furthermore, comparing two cavity types (natural and human-made), we demonstrated that artificial nestboxes, a tool widely used in field ecology, can impact the microbiome assembly in nestlings. Our results align with and expand upon earlier insights in great tit *Parus major* gut microbiome variation inferred in the same urban study system on a species ecologically related to the blue tit, confirming the pervasive impact of urbanisation on the avian gut microbiome.

## Introduction

Urbanisation creates a novel ecosystem with a set of specific conditions that pose an adaptive challenge for the wild animals inhabiting it - this includes, among many, habitat fragmentation, warmer temperatures, altered resource availability, higher population densities, and exposure to human disturbance (Garrard et al., 2017; Douglas et al., 2010; Szulkin et al., 2020). While these environmental changes may benefit certain species, others face significant challenges, leading to reduced survival, reproductive success, and overall fitness (Lowry et al., 2013).

For example, urban birds tend to lay their eggs earlier in the season and have smaller clutch sizes compared to their rural counterparts (Chamberlain et al., 2009, Capilla-Lasheras et al., 2022). Both juvenile and adult birds from urban environments also tend to be smaller and have a lower body mass (Chamberlain et al., 2009; Corsini et al., 2021; Thompson et al., 2022), and while the stable availability of anthropogenic food positively influences adult well-being, the paucity of natural food sources poses a threat to the nourishment of nestlings (Chamberlain et al., 2009). Findings stemming from study sites located in Warsaw, Poland, where our study took place, are consistent with these general patterns: the percentage of impervious surface area around nestboxes was shown to have a negative impact on offspring development, body mass, and survival in both blue tits and great tits (Corsini et al., 2021; Corsini & Szulkin, 2025).

The gut microbiome is an important fitness component as it influences several aspects of host biology (Grond et al., 2018), and the bird microbial assemblage patterns may be shaped by both intrinsic and extrinsic factors, including social relationships, diet, and environmental conditions (Sun et al., 2022). Consequently, urbanisation can impact the gut microbiome through various mechanisms, making it a focal point of interest for urban ecologists. For example, in American white ibises lower microbial diversity was correlated with the quantity of digested human-provisioned food (Murray et al., 2020). In a study conducted in Kenya on 57 avian species, it was found that increasing human density, coupled with higher livestock density, may be linked to the spillover of antimicrobial resistance genes into birds, potentially facilitating the spread of resistant pathogens and impacting both wildlife and public health (Hassell et al., 2019). A similar investigation involving samples from 30 different bird species collected in eight countries revealed that the prevalence of antimicrobial resistance genes in gut microbiomes was six times higher in urban environments (Mourkas et al., 2023). In a study on house sparrows urbanisation was associated with a lower taxonomic diversity of microbiome and changes in community structure (Teyssier et al., 2018). Interestingly, a study on the same study system reported a substantial impact of the availability of anthropogenic food sources on glucose levels and decreased uric acid levels in the blood - both of which could potentially influence microbiome composition (Gadau et al., 2019). Contrary to the notion of lower diversity being a prevailing characteristic of urban microbiomes, urban birds in a study on white-crowned sparrows exhibited higher microbial diversity. This outcome, however, was associated with more variable land cover types in the urban areas under investigation compared to non-urban areas, and the elevated microbial diversity was correlated with a higher tree cover density (Phillips et al., 2018). This result underscores the significance of the local environment, challenging the binary urban and rural dichotomy that inaccurately lumps diverse environments such as woodlands and office areas based on exposure to disturbances (Maraci et al., 2022).

Overall, studies on wild microbiomes in urban environments reveal patterns that are often inconsistent and highly variable, mainly due to the lack of spatial and temporal replication. As the sensitivity to disturbances may be species-dependent, it is also crucial to examine the differences between various species sharing the same habitat. We additionally remark on a visible need for exact quantification of land use that would allow between-location comparisons. Moreover, microbiome studies have predominantly focused on adult birds, and little attention has been given to juveniles. It is thus essential, considering the current scale of anthropogenic pressures, to better understand how resilience to disruptions may be shaped in the earliest stages of development.

Until recently, the only study exploring this aspect in the urban space was conducted on great tits, using the same gradient of urbanisation and methodology as described here. The results revealed that nestlings growing up in more highly urbanised environments exhibited lower gut microbial diversity (Maraci et al., 2022). Building upon these observations and aiming to fill these knowledge gaps, we focused on blue tit (*Cyanistes caeruleus*), an avian species not previously examined in terms of its gut microbial diversity in an urban setting, specifically at a life stage when the microbiome is still under development. Using 16S rRNA gene sequencing, we investigated the gut microbiome of blue tits from an urban mosaic of a European capital city, where we characterised heterogeneity between microhabitats, in 2018 and 2019. We tested whether (i) urbanisation impacts individual microbial diversity and composition of blue tit nestlings; (ii) these patterns are repeatable across years; and (iii) nestboxes, as examples of human-made structures, alter the microbial diversity when compared to natural cavities. Additionally, we aimed to assess how generalisable or different these patterns are between two species, great tits and blue tits, by comparing the findings of the present study with those produced for great tits (both species were sampled in the same study locations in the same year).

## Results

Faecal samples from 15-day-old blue tit nestlings were collected in 2018 and 2019 as part of a long-term monitoring effort of the breeding activity of blue tits and great tits in a gradient of urbanisation in Warsaw, Poland. We analysed a total of 107 samples originating from 91 nestboxes (Schwegler, type 1b) distributed across 9 study sites located within and outside of the city (urbanisation gradient) and from 16 natural cavities located in an urban forest. Given that environmental conditions, including weather patterns and food availability, can vary between years and influence avian microbiome, we examined whether observed patterns were consistent across both breeding seasons. To focus on the impact of urbanisation in nestlings reared in the same cavity type, the basic diversity and community structure analyses solely used data derived from nest boxes in the urbanisation gradient. To compare the impact of cavity type on microbiome, we used samples collected across one, highly homogenous location: the urban Bielany forest, where nests in natural cavities (i.e. holes in trees) and artificial cavities (i.e. nestboxes) were present (Supplementary Table 3).

### Microbial community diversity and composition in relation to urbanisation and year of sampling

We examined alpha diversity in relation to urbanisation with linear mixed models to account for temporal variation (two years of sampling and differences in individual hatching dates), spatial structure (percentage of Impervious Surface Area as a measure of urbanisation, and sampling site assignment), and potential survival-related health differences (fledging - successful or not). We used three key diversity metrics as response variables in separate models: Shannon’s index, Chao1, and Faith’s phylogenetic diversity. The model with Shannon diversity as the response variable was run without the ISA*year interaction as it was not statistically significant. We observed an overall lower alpha diversity in nestboxes characterised by high ISA percentages in their immediate environment (i.e. more urbanised) compared to nestboxes located in low ISA percentage areas (i.e. less urbanised) (Table 1). Increasing values of ISA thus covaried with lower Chao1 and Faith’s PD indices, revealing that alpha diversity decreased with increasing amount of impervious surfaces around the nestbox (Table 1). This effect, however, was year-dependent: in 2018, alpha diversity decreased with increasing ISA, while no such relationship was observed in 2019 (Table 1, Figure 1).

**Table 1:** Variation in gut microbiome alpha diversity as a function of Impervious Surface Area, year and their interaction, sample collection day, and whether or not the nestling survived until fledging. Significant results are indicated in bold.

| Fixed effect | Response variable | Est | CI | p |
| --- | --- | --- | --- | --- |
| ISA * year | **Chao1** | **0.03** | **0.00 - 0.06** | **0.041** |
|  | **Faith's PD** | **0.02** | **0.00 - 0.04** | **0.042** |
| ISA | **Chao1** | **-0.02** | **-0.04 - -0.00** | **0.044** |
|  | Shannon | -0.00 | -0.01 - 0.01 | 0.795 |
|  | **Faith's PD** | **-0.02** | **-0.03 - -0.01** | **0.008** |
| year [2019] | **Chao1** | **-0.7** | **-1.2 - -0.2** | **0.006** |
|  | Shannon | -0.21 | -0.32 - 0.05 | 0.162 |
|  | **Faith's PD** | **-0.43** | **-0.78 - -0.08** | **0.016** |
| sample collection day | Chao1 | -0.07 | -0.35 - 0.21 | 0.619 |
|  | Shannon | -0.09 | -0.19 - 0.05 | 0.259 |
|  | Faith's PD | -0.12 | -0.32 - 0.08 | 0.242 |
| survival | Chao1 | 0.44 | -0.22 - 1.09 | 0.186 |
|  | Shannon | 0.09 | -0.16 - 0.39 | 0.399 |
|  | Faith's PD | 0.38 | -0.08 - 0.84 | 0.103 |

**Figure 1:**
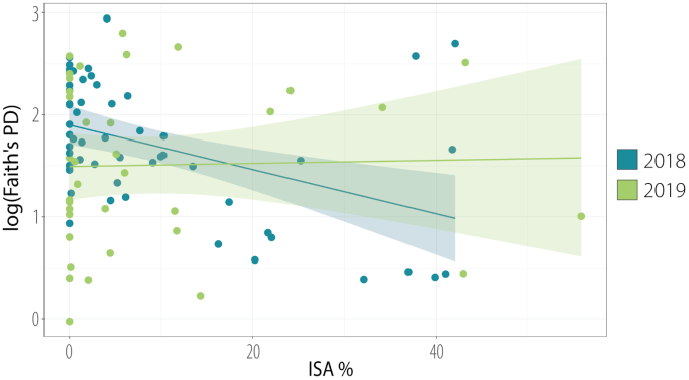
Year-dependent alpha diversity patterns in a gradient of urbanisation. The regression lines illustrate the covariation between the percentage of Impervious Surface Area and Faith’s Phylogenetic Diversity index (logarithmic scale).

In all models of beta diversity (PERMANOVA with four dissimilarity matrices: Jaccard, Bray-Curtis, UniFrac, and Weighted UniFrac), microbial composition varied between years (Table 2; Figure 2). All matrices but Weighted Unifrac also revealed differences related to the percentage of Impervious Surface Area and the site of sampling (Table 2). To examine whether this result was driven by sampling year, we ran separate models for each year and, in accordance with the results for alpha diversity (Figure 1), the impact of urbanisation on beta diversity was visible in 2018, whilst there was no such effect in 2019 (Table 3). The analysis also uncovered differences driven by the phenology of sampling in the season in Bray-Curtis and UniFrac matrices (Table 2).

**Table 2:**
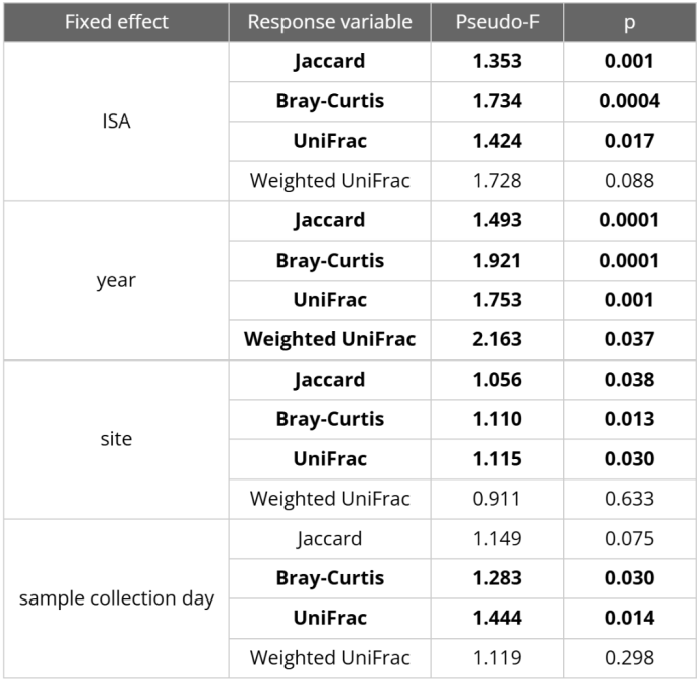
Compositional differences in response to different Impervious Surface Area percentages, years, sampling sites, and sample collection dates were tested using PERMANOVA models. Significant results are indicated in bold.

| Fixed effect | Response variable | Pseudo-F | p |
| --- | --- | --- | --- |
| ISA | **Jaccard** | **1.353** | **0.001** |
|  | **Bray-Curtis** | **1.734** | **0.0004** |
|  | **UniFrac** | **1.424** | **0.017** |
|  | Weighted UniFrac | 1.728 | 0.088 |
| year | **Jaccard** | **1.493** | **0.0001** |
|  | **Bray-Curtis** | **1.921** | **0.0001** |
|  | **UniFrac** | **1.753** | **0.001** |
|  | **Weighted UniFrac** | **2.163** | **0.037** |
| site | **Jaccard** | **1.056** | **0.038** |
|  | **Bray-Curtis** | **1.110** | **0.013** |
|  | **UniFrac** | **1.115** | **0.030** |
|  | Weighted UniFrac | 0.911 | 0.633 |
| sample collection day | Jaccard | 1.149 | 0.075 |
|  | **Bray-Curtis** | **1.283** | **0.030** |
|  | **UniFrac** | **1.444** | **0.014** |
|  | Weighted UniFrac | 1.119 | 0.298 |

**Table 3:**
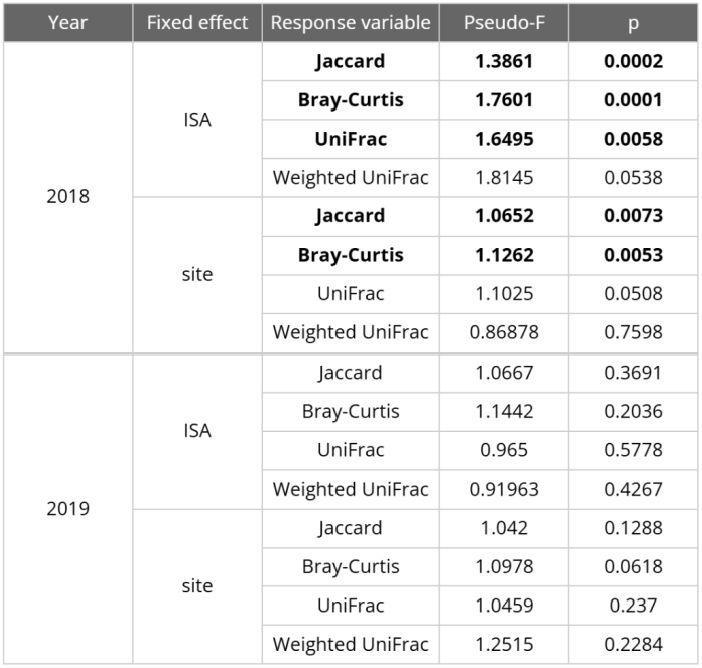
Results of PERMANOVA analyses for 2018 and 2019 datasets. The significant results are indicated in bold, revealing year-specific effects: Impervious Surface Area and sampling sites did influence the compositional similarity of microbiomes, but only in 2018.

**Figure 2:**
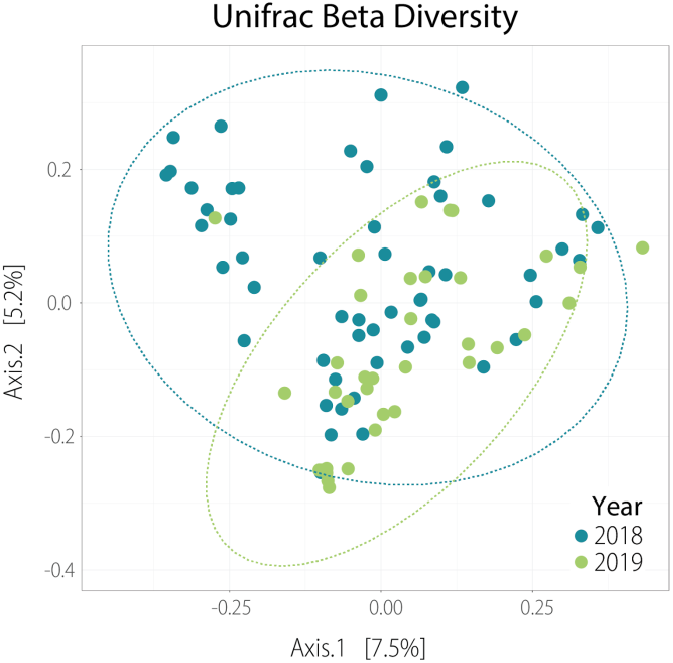
UniFrac Beta matrix illustrating dissimilarities in the compositional structure of samples collected in the years 2018 and 2019.

The homogeneity of multivariate dispersions was additionally tested between year groups with PERMDISP. The median distances for UniFrac beta diversity, but for this matrix only, were not even (p = 0.01228), suggesting that the PERMANOVA results, in this case, can be impacted by different levels of internal variability of beta diversity within year groups. In order to account for the significant variation between years in beta diversity, differential abundance analysis was applied, revealing higher abundances of *Streptococcaceae* and unclassified bacteria from the order Chlamydiales in 2018 (Supplementary Table 1).

### Impact of cavity type on nestlings microbiome

To analyse the impact of cavity type on nestling microbiome, we generated a subset of samples collected from natural cavities and nestboxes located in the same urban forest across two years (in total, 16 samples from natural cavities and 17 samples from nestboxes). We did not include nestboxes from locations other than the urban forest to eliminate the possible influence of any varying environmental conditions, especially to exclude differences in urbanisation pressure.

Taking into account the possible impact of year, we ran models that included the interaction between year and cavity type. Shannon index, being one of the three indices that we used, revealed significantly lower microbial diversity in samples collected from natural cavities and a strong effect of the interaction between year and cavity type (Table 4; the remaining models in Supplementary Table 2a). To better understand how cavity type and year interact, we ran post hoc tests, which revealed different year-to-year changes in nestboxes and natural cavities: in nestboxes, in 2019 the microbial diversity was lower than in 2018 (Shannon: p = 0.0118; Figure 3) while in samples from natural cavities, gut microbiome alpha diversity remained stable between years (Shannon: p = 0.979; Figure 3; Supplementary Table 2b).

**Table 4:** Variation in gut microbiome alpha diversity (Shannon index) as a function of year, cavity type, and their interaction. Reference categories are 2018 for the year and natural cavity for cavity type. Diversity was significantly lower in 2019 and in natural cavities. Significant results are indicated in bold.

| Predictors | Est | CI | p |
| --- | --- | --- | --- |
| (Intercept) (2018, natural cavity) | 3.24 | 2.6 - 3.89 | <0.001 |
| year (2019 vs. 2018) | 1.94 | 1.13 - 2.76 | 0.699 |
| cavity type (nestbox vs. natural) | **1.3** | **0.26 - 2.34** | **0.016** |
| year * cavity type | **2.38** | **-4.13 - -0.62** | **0.01** |

**Figure 3:**
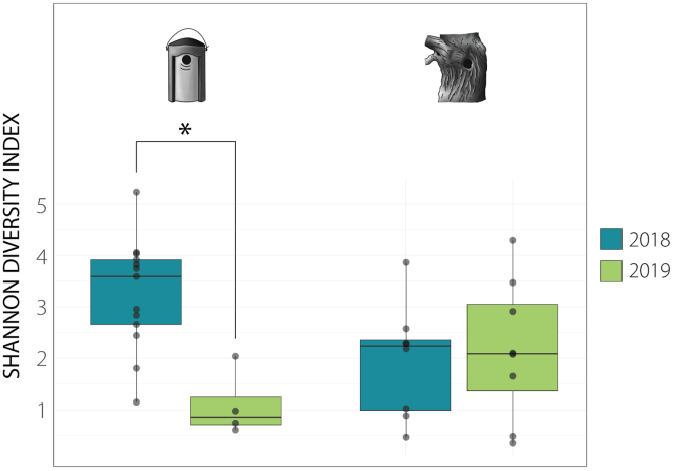
Shannon diversity index boxplots (with medians, 25th and 75th percentiles, respectively) for nestboxes and natural cavities across years. Differences between the years are shown in colour. The significant differences revealed by the post hoc test were determined based on the linear model at p values ≤ 0.05 (*), p ≤ 0.01 (**) and p ≤ 0.001 (***).

None of the PERMANOVA analyses used to test for beta-diversity dissimilarities in composition between years and cavity type revealed any differences - the exception was the effect of year in the Bray-Curtis matrix (p = 0.0377; Supplementary Table 2c).

## Discussion

In our study, we explored the impact of urbanisation variability on the gut microbiome of blue tit nestlings over two years. Additionally, we tested the impact of cavity type (natural vs. human-made). We measured urbanisation with a continuous value of ISA and noted that it negatively impacts individual microbial diversity and greatly influences the compositional similarity of microbiomes in comparably urbanised areas. The obtained results highlight the importance of temporal replication of ecological studies, remarking that the pressure the urban environment exerts on avian microbiome can vary over time - in our case, this variation was clearly visible when samples were collected in just two contrasting years. Moreover, the comparison between natural and artificial cavities suggests that these two cavity types provide immediate environments that may differently support microbial assembly in the first days of nestling life, in line with earlier findings in great tits in the same study system (Maraci et al. 2022).

### The degree of urbanisation can be reflected by the similarity of microbial composition and its diversity

Our results build upon an earlier study on great tits (Maraci et al., 2022), which took place in one of the same years (2018) as the current study, and in the exact same locations. Importantly, Maraci et al. (2022) also reported a negative, significant correlation between ISA and microbial alpha diversity. This suggests that these two closely related species that inhabit the same niches might respond similarly to environmental stimuli. Studies on other species investigating how microbial alpha diversity varies in urban spaces have revealed mixed results. Similarly to the results reported in Warsaw, lower diversity in urban areas was observed in adult house sparrows (Teyssier et al., 2018; Teyssier et al., 2020) and herring gulls (Fuirst et al., 2018). Conversely, gut microbial diversity was higher in urban populations of white-crowned sparrows than in their rural counterparts (Phillips et al., 2018), and no significant results were found in American white ibises (Murray et al., 2020).

The discrepancies between studies may result from differently defined urbanisation measures (urban/rural, percentage of built-up area, human population, or tree cover density) or simply originate from taxonomy or life-stage history. To capture possible patterns, we would need to use a standardised measuring system that does not raise any doubts regarding the definition of urbanisation of the area (Maraci et al., 2022) and replicate the studies on the same species to observe whether emerging trends are related to the taxonomy or rather the particularity of the studied environment (Szulkin et al., 2020). Additionally, a meta-analytical quantification of effect sizes across studies could be performed to assess whether trends are emerging across species.

Although some studies showed associations between higher microbial diversity and good health (reviewed in Ottinger et al., 2024), it is known that microbiome tends to fluctuate in early life (Maraci et al., 2022; Somers et al., 2023). The fitness consequences of these variations in gut microbiome should be further investigated. To gain better insight into the possible implications of differences in microbial diversity, it would be necessary to assess not only species diversity but also their functional diversity (Worsley et al., 2021).

### Year-specific effects of urbanisation on microbial diversity

The analysis of samples collected over two years revealed highly significant temporal differences in gut microbiome variation, suggesting that the impact of urbanisation on developing nestlings can be highly year-specific. Since sampling site characteristics remained the same in 2018 and 2019, the most likely drivers for observed microbiome alterations are differences in climatic conditions regarding temperature, rainfall, humidity, and the downstream differences related to food availability. Indeed, as described by Sudyka et al. (2022) in a study conducted on blue tits in the same years and in one of the exact locations as the data collected here (urban forest), weather conditions differed markedly between these years (Sudyka et al., 2022). Weather data from a local weather station indicated that in 2019, April and May were significantly colder in terms of temperature average, minimum and maximum. Additionally, in 2019, April and June tended to be drier, May was more humid with higher precipitation, and the entire season was windier. All these conditions suggest that 2019 was a less favourable year. Importantly, these climatic alterations can directly influence the availability of primary food sources such as caterpillars and other invertebrates (Visser et al., 2006). As the type and amount of food influence the gut microbiome (Xiao et al., 2021; Schmiedová et al., 2022), the observed differences between years can be attributable to the direct effect of food availability on the gut microbiome. Furthermore, these months are crucial for nestling growth, and changes in food availability might lead to differential growth rates (Corsini et al., 2021), ultimately influencing microbial assembly. The increasing unpredictability of weather patterns, as a result of the human-induced global climate crisis, may drastically interfere with ecosystem stability and resilience (Malhi et al., 2020). In environments subjected to frequent and rapid weather fluctuations, the unpredictability of foraging and feeding activities generates irreparable consequences on the endocrine and metabolic systems of both nestlings and their parents (Fokidis et al., 2012). This could also be reflected in the microbiome.

The two families detected in higher abundances in 2018, Streptococcaceae and unclassified Chlamydiales (Supplementary Table 1), are known to encompass some pathogenic bacterial species, and therefore their increased abundance might pose a health risk. Given that different weather conditions led to varying food availability between the years, these may have also contributed to differences in the bacterial abundances - certain microbial families may, in fact, thrive in response to specific dietary components (Teyssier et al., 2020). Although some genera included in these families are commonly known as human pathogens, beneficial interactions have also been observed in wild animals. For instance, higher levels of *Streptococcaceae* were detected in healthy wild mallards compared to mallards infected by influenza A viruses (Ganz et al., 2017). Chlamydiales, on the other hand, are more frequently associated with bacterial infections in wild birds and can spill over to domestic animals, posing risks to public health (Burnard & Polkinghorne, 2016).

### Differential impact of natural cavities and nestboxes on nestling microbial diversity

By using samples collected only in a single location (urban forest), we eliminated the possible impact of varying levels of urbanisation and other external factors such as food availability. This allowed us to specifically test the effect of cavity type. Nestboxes, most frequently used in studies on cavity-nesting birds, differ on many grounds from the natural nesting environment - most notably, providing poorer insulation, leading to lower humidity and higher fluctuations in daily ambient temperatures (Maziarz et al., 2017; Sudyka et al., 2023). These differences can impact various aspects of early development. For example, the study by Sudyka et al. (2022), conducted on blue tits in the same years and location as the data collected for this study, focused on reproductive success dependent on cavity type. Blue tits breeding in nestboxes produced fewer nestlings and had lower hatching and fledging success compared to those nesting in natural cavities.

Results reported here show that alpha diversity was dependent on cavity type, with the microbiome being generally more diverse in nestboxes. However, it is essential to remark that these results were again year-dependent: while in 2018, the median alpha diversity was higher in nestboxes (importantly, the same effect was observed in our previous study on great tits where samples were collected only in 2018; Maraci et al., 2022), this trend reversed in 2019. Strikingly, the year-specific patterns differed between the cavity types: in nestboxes, microbial diversity dropped significantly in 2019, while in natural cavities, diversity remained at comparable levels between years (Figure 3). This result could possibly imply higher susceptibility of nestlings reared in nestboxes to external conditions. The aforementioned study on blue tit breeding success (Sudyka et al. 2022) also reported a significantly longer duration of nesting in birds raised in nestboxes. This could suggest that even though all of them were examined and sampled on their 15th day of life, their microbial communities might have been at different stages of development. Additionally, their microbiomes could have been influenced by the overall higher temperature and its fluctuations in nestboxes (Sudyka et al., 2022), which could mediate immune and metabolic functions, although the relationship between temperature and microbial community assembly remains mostly unknown in wild passerine birds. The only study up to date that has touched upon this aspect, performed on great tits *Parus major*, found an association between lower temperatures and higher alpha diversity (Liukkonen et al., 2024).

### Conclusions and outlook

Our study serves as an extended continuation of previous observations of gut microbiome variation in great tits (Figure 4), which we have referred to multiple times throughout the text (Maraci et al., 2022). We report a highly equivalent pattern of decreasing alpha diversity with increasing urbanisation, as observed in the same year (2018) and study system. In terms of beta diversity in blue tits, community composition differed with changing ISA percentages. In great tits, no such relationship between ISA and composition was detected, although there were marked differences in microbial composition between urban and rural areas. This might indicate species-specific differences in the sensitivity of microbial composition to fine-scale alteration in the local environment. Additionally, in both species, the type of cavity impacted alpha diversity, with lower values observed in natural cavities. In summary, these results highlight the consistency and complementarity of our studies, providing deeper insight into urban-driven relationships between the two species that share the same ecological niche and parallel gut microbiome variation.

**Figure 4:**
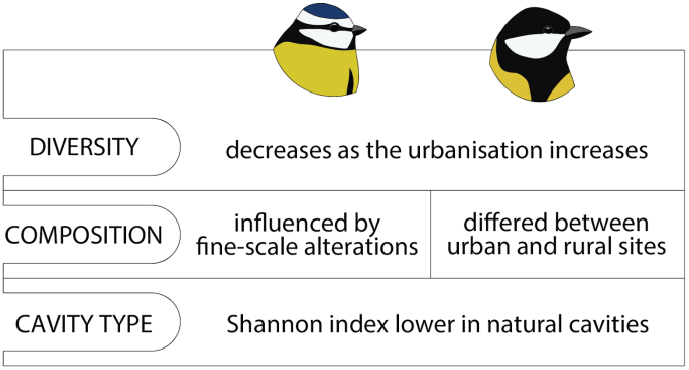
Comparison of results obtained in this study and in the previous study on great tits (Maraci et al., 2022). The microbial alpha diversity of both species in 2018 was negatively correlated with urbanisation. Alpha diversity (Shannon index) was also lower in natural cavities than in nest boxes. However, the impact of urbanisation on the structural similarity of microbiomes (beta diversity) was species-specific.

All in all, this study showed clear urban-related alterations in the gut microbiome of a small passerine. However, these results were also found to be year-dependent, highlighting the critical importance of study replication in terms of avoiding overgeneralisation. Additionally, we observed higher diversity in microbiomes of nestlings originating from nestboxes, but their microbiomes were notably more susceptible to change between years than those from natural cavities. While providing valuable insights into how the avian microbiome responds to anthropogenic alteration, our study can also benefit from further refinements: it would be valuable to investigate urban microbiome variation through taxonomic, spatial and temporal replication. A valuable follow-up includes investigating how urban diet and food availability shape microbiome composition, as well as how it can possibly be impacted by weather conditions, particularly in the context of their globally observed unpredictability. Answering these questions would allow us to better comprehend the ecological and evolutionary response of non-human animals to human-induced rapid anthropogenic change. With rapidly increasing global climate change and urbanisation and the progressing disruption of natural habitats, we emphasize the importance of examining the microscale of organismal dependencies to understand how our welfare depends on the seemingly invisible interconnectedness between us and other beings.

## Methods

### Study sites and environmental metrics

To better reflect urban landscape heterogeneity, the sampling locations, especially within the city borders, varied greatly in the environmental structuring: they spanned from natural and urban forests, through peri-urban village and urban woodlands, to residential and office areas (Figure 5). Therefore, we decided not to assign the dichotomous categories of urban and rural and instead used a more precise urbanisation measure with values specific to each individual sampling location. We used the percentage of Impervious Surface Area (referred to as ISA) around each nest location (measured in 100 m radius around each nestbox in the study system, and calculated using the imperviousness map downloaded from Copernicus Land Monitoring Services (Szulkin et al., 2020). The 100 m radius was used based on the food foraging distance observed in this species while feeding the nestlings (Tremblay et al., 2004). ISA includes both all built-up areas and soil-sealing surfaces, and strongly covaries with various environmental parameters (such as tree cover density; Szulkin et al., 2020) that may influence the gut microbiome by impacting food availability and, consequently, diet composition. All of the environmental parameters have been calculated for each nestbox as part of the long-term monitoring effort, and they included spatial variables (distance to the city centre, closest road, and closest path), variables collected on the ground (human presence, sound pollution, and temperature), and variables extrapolated from digital photography and satellite imagery (tree cover density and light pollution) (in detail in Supplementary Text 1).

**Figure 5:**
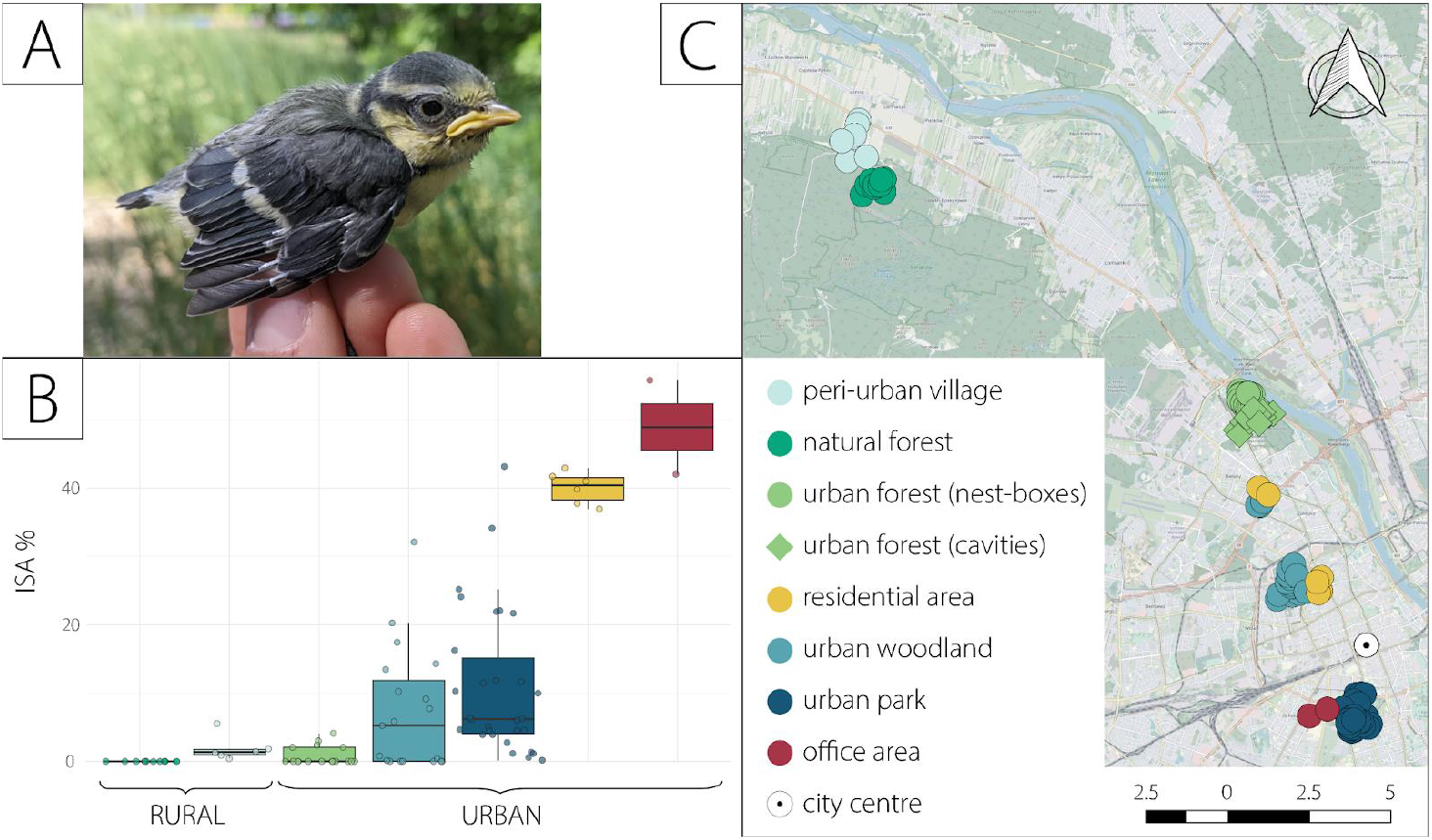
a) Study species: 15 day old blue tit *Cyanistes caeruleus* on sampling day; b) Boxplot illustrating proportions of samples in every given location with marked median values of Impervious Surface Area percentage; c) Map of sampling locations.

### Data collection and laboratory work

Sample collection took place depending on individual hatching dates and spanned from May 16 to June 10 in 2018 and May 12 to June 23 in 2019. As food availability may change with time, and diet constitutes an important aspect of microbial community structure, the temporal variability was taken into account by including standardised sampling date. The Julian date of sampling (15th day of life, dates varied for individual nestlings) was standardised by subtracting the population sampling mean and dividing it by its standard deviation, obtaining a population-wide mean sampling date of zero and a standard deviation of 1 for every study year. Faeces were collected from one chick per nest using non-invasive methods and deposited directly in 5 ml sterile Eppendorf tubes filled with 3 ml of RNAlater (Qiagen), and stored at −20°C and transferred to the University of Bielefeld in Germany.

The laboratory work was carried out in accordance with the pipeline previously described in the earlier study on great tit nestling microbiome, which was performed on the same urbanisation gradient (Maraci et al., 2022). Microbial DNA was extracted using the DNeasy PowerLyzer PowerSoil Kit (Qiagen), as described in the manufacturer’s protocol. The hypervariable V3-V4 region of the 16S ribosomal RNA gene was targeted following Illumina 16S Metagenomic Library Preparation Guide, 15044223-B. The final amplicon pool, alongside 119 biological samples, contained two blank controls for DNA extraction and one blank control for PCR amplification. All of these were sequenced on the Illumina MiSeq system (Illumina, Inc., San Diego, CA, USA) in CeBiTec at the University of Bielefeld.

### Data analysis

The bioinformatic processing was performed using the nf-core/ampliseq standardised pipeline (Straub et al., 2020) and involved subsequent steps. First, the raw sequencing data in FASTQ format was subjected to FastQC and the overall quality of reads was assessed manually. Then, the amplification primers (FW-5’-CCTACGGGNGGCWGCAG-3’ and RV-5’-GACTACHVGGGTATCTAATCC-3’) were used to trim the reads using Cutadapt (Martin, 2011). This process required a minimum primer overlap of ten bases with a maximum error rate of 10%. The remaining pool was quality filtered and denoised using DADA2 (Callahan et al., 2016) to correct sequencing errors, remove chimaeras, and create the output of amplicon sequence variants (ASVs). ASVs were then filtered for ribosomal RNA sequences only, using Barrnap (Seemann, 2013) as a prediction tool. Eventually, taxonomic classification of the ASVs was assigned using DADA2 and SILVA v138 as a reference database (Pruesse et al., 2007). The phylogenetic tree was generated using Pplacer (Matsen et al., 2010).

All of the following statistical analyses were conducted in R version 4.3.1 (R Core Team, 2021) and PRIMER v7 (Clarke & Gorley, 2015). The raw dataset was initially filtered for contaminants with the help of decontam R package (Davis et al., 2018) - in total, 42 potential contaminant ASVs were excluded from the dataset. In order to account for the uneven distribution of the number of reads between samples (min = 1136, max = 46033), all samples that remained after filtering were rarefied to a depth of the lowest read count observed, leaving 3488 ASVs across 107 samples (restricted from 119 as a result of quality filtering).

The impact of urbanisation was measured using three different metrics of alpha diversity: Shannon’s diversity index (phyloseq R package, 1.46.0) (McMurdie & Holmes, 2013), Chao1 (phyloseq R package, 1.46.0) (McMurdie & Holmes, 2013), and Faith’s phylogenetic diversity (picante R package, 1.8.2) (Kembel et al., 2010). These metrics of microbiome diversity were then assessed in the context of impervious surface variation estimated around each nestbox.

Due to the non-normal distribution of values in each of these diversity indices, the data was normalised employing square root transformation in the case of Shannon diversity and logarithmic transformation in the remaining two. Three linear mixed models were fitted using the lme4 R package (version 1.1-34; Bates et al., 2015), using ISA, year and its interaction as fixed effects, and the diversity metrics as the response variables. The interaction aimed to test whether a possible ISA effect on microbiome diversity was year-specific. To account for the potential impact of differences in the sample collection dates on alpha and beta diversity, the number of days between the initiation of the study and sample collection was added as a fixed effect. Additionally, the survival of nestlings was inferred as a fixed effect in alpha diversity analysis (11 samples came from nestlings that did not fledge from the nestboxes, which could suggest their impoverished health reflected in the microbial diversity). The associations between sampling sites were also added to the model as a random variable to account for the non-independence of the samples originating from the same sampling site. The spatial autocorrelation was also tested in the simulated scaled residuals of the fitted LMMs using Moran’s I test with the use of DHARMa package (version 0.4.6; Hartig, 2018).

To test whether the microbial communities were affected by cavity type, we employed a subset of data, including samples collected only from the urban forest (17 samples from nestboxes and 16 from natural cavities, one sample per nest). Chao1 and Faith’s PD values were normalised employing logarithmic transformation, and three basic linear models that included interaction between the cavity type and year were fitted.

To visualise the between-group differences in community composition, Principal Coordinate Analysis plots were generated for four types of dissimilarity matrices: Jaccard (quantitative, prevalence-based; Jaccard, 1912), Bray-Curtis (quantitative, with bacterial abundances; Bray and Curtis, 1957), Unifrac (qualitative, phylogenetic, prevalence-based; Lozupone and Knight, 2005), and Weighted Unifrac (which also accounts for abundances; Lozupone et al., 2007). Then, PERMANOVA models were fitted, taking into account the ISA values, year, site, and sampling date. The models for the subset, including natural cavities focused on cavity type and year.

Finally, to detect finer differences between years and to determine which ASVs were differently abundant, we employed corncob analysis (0.4.1; Martin et al., 2020). To detect ecologically relevant patterns, we focused on the family level.

## Ethics statement

The research was carried out with a permit from the Regional Directorate for Environmental Protection (RDOŚ) in Warsaw, Poland.

## Data accessibility

The scripts used for processing the data are provided in the GitHub repository at: https://github.com/lena-fus/blue-tits-warsaw

## Author contributions

L.F., Ö. M., and M. S. conceptualized the research idea and planned the experiments. M. S., I. dL. and J. S. collected the biological samples and environmental data. L.F. and Ö. M. carried out the laboratory experiments. S. J. performed the bioinformatic analyses. L.F. carried out the statistical analyses with the supervision of Ö. M. and M.S., with input from J.S. L.F. wrote the manuscript in consultation with Ö. M and M. S., and all authors approved the final version of the manuscript.

## Acknowledgements

We are grateful to all members of the Anthropocene Biology Lab for collecting data in Warsaw and its surrounding regions. This paper was typeset with the bioRxiv word template by @Chrelli: www.github.com/chrelli/bioRxiv-word-template

