## Supplementary material for "Gut microbiome variation in juvenile blue tits in a European urban mosaic"

**Supplementary Table 1.** Results of differential abundance of microbial families between years. Both listed families were more abundant in 2018.

| padj | p | Phylum | Family |
|---:|---:|---|---|
| 0.0084594 | 0.0000379 | Verrucomicrobiota | Chlamydiales unclassified |
| 0.0212351 | 0.0001904 | Firmicutes | *Streptococcaceae* |

**Supplementary Table 2a.** Linear models: alpha diversity between two types of cavities and two years.

| Predictors | Response variable | Est | CI | p |
|:---:|---|---:|---:|---:|
| year (2019 vs. 2018) | Chao1 | -13.39 | -92.19 - 65.41 | 0.731 |
| | Shannon | 0.22 | -0.93 - 1.38 | 0.699 |
| | Faith's PD | -0.79 | -5.09 - 3.5 | 0.709 |
| cavity type (nestbox vs. natural) | Chao1 | 40.97 | -29.85 - 111.78 | 0.246 |
| | **Shannon** | **1.3** | **0.26 - 2.34** | **0.016** |
| | Faith's PD | 2.65 | -1.21 - 6.51 | 0.171 |
| year * cavity type | Chao1 | -81.72 | -201.43 - 37.98 | 0.173 |
| | **Shannon** | **-2.38** | **-4.13 - -0.62** | **0.01** |
| | Faith's PD | -5.66 | -12.18 - 0.87 | 0.087 |

**Supplementary Table 2b.** Results of post-hoc tests (year and cavity type).

| Contrast | | Index | Est | SE | p |
|---|---|---|---|---|---|
| BOX-2018 | NAT-2018 | Shannon | 1.301 | 0.507 | 0.071 |
| | | Faith's PD | 0.4475 | 0.284 | 0.4075 |
| **BOX-2018** | **BOX-2019** | **Shannon** | **2.156** | **0.646** | **0.0118** |
| | | **Faith's PD** | **1.0972** | **0.361** | **0.0244** |
| BOX-2018 | NAT-2019 | Shannon | 1.08 | 0.507 | 0.1681 |
| | | Faith's PD | 0.5221 | 0.284 | 0.2763 |
| NAT-2018 | BOX-2019 | Shannon | 0.855 | 0.692 | 0.6096 |
| | | Faith's PD | 0.6497 | 0.387 | 0.3526 |
| NAT-2018 | NAT-2019 | Shannon | -0.221 | 0.565 | 0.9794 |
| | | Faith's PD | 0.0746 | 0.316 | 0.9953 |
| BOX-2019 | NAT-2019 | Shannon | -1.075 | 0.692 | 0.4192 |
| | | Faith's PD | -0.5751 | 0.387 | 0.4584 |

**Supplementary Table 2c.** Results of PERMANOVA tests for beta diversity between cavity types.

| Fixed effect | Response variable | Pseudo-F | p |
|---|---|---|---|
| year | Jaccard | 1.11781 | 0.0522 |
| | **Bray-Curtis** | **1.2337** | **0.0377** |
| | UniFrac | 1.15088 | 0.1321 |
| | Weighted UniFrac | 1.0815 | 0.3145 |
| cavity type | Jaccard | 1.0598 | 0.1642 |
| | Bray-Curtis | 1.0923 | 0.1867 |
| | UniFrac | 1.15587 | 0.1345 |
| | Weighted UniFrac | 1.1812 | 0.2491 |

**Supplementary Table 3.** Sample size and the distribution of ISA among the sampling sites.

| Habitat type | Cavity type | Initial N | Final N | Min | Max | Mean | SD | Median |
|---|---|---|---|---|---|---|---|---|
| peri-urban village | nest box | 6 | 6 | 0.41 | 5.53 | 1.89 | 1.85 | 1.36 |
| natural forest | nest box | 12 | 11 | 0.00 | 0.00 | 0.00 | 0.00 | 0.00 |
| urban forest | nest box | 19 | 17 | 0.00 | 4.09 | 0.80 | 1.35 | 0.00 |
| | natural cavity | 17 | 16 | NA | NA | NA | NA | NA |
| residential area I | nest box | 4 | 4 | 36.97 | 42.95 | 40.38 | 2.60 | 40.79 |
| residential area II | nest box | 3 | 2 | 37.74 | 41.00 | 39.37 | 2.30 | 39.37 |
| urban woodland I | nest box | 17 | 16 | 0.00 | 32.13 | 5.89 | 8.99 | 0.60 |
| urban woodland II | nest box | 3 | 3 | 9.11 | 20.25 | 14.27 | 5.61 | 13.46 |
| urban park | nest box | 36 | 30 | 0.16 | 43.16 | 10.89 | 10.68 | 6.20 |
| office area | nest box | 2 | 2 | 42.04 | 55.81 | 48.92 | 9.73 | 48.92 |
| | *total:* | 119 | **107** | | | | | |

**Supplementary Figure 1.** Pearson's correlation coefficients among ISA and environmental variables.

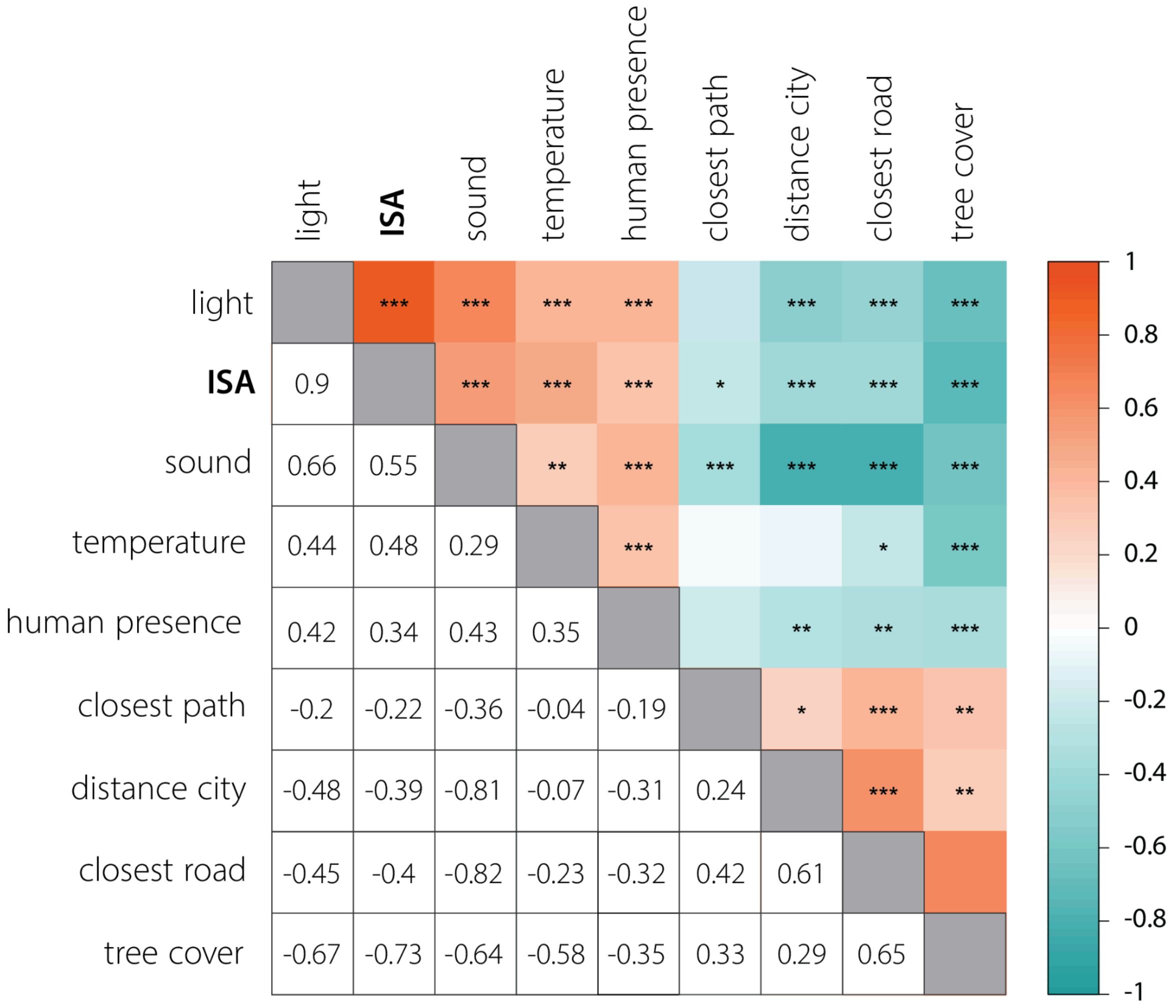

**Supplementary Text 1.** Environmental and spatial variables used in the study.

1. **variables collected on the ground:**

    (a) human presence, derived by quantifying all humans and dogs within a 15 m radius around each nest box (Corsini et al., 2017),

    (b) sound pollution, obtained after averaging recordings on the DbC scale using hand-held sound level metres over four days throughout the field season (Szulkin et al., 2020), and

    (c) temperature, measured with Thermocrone ibuttons DS1921G set in 2018 from April 24 until June 30 (Szulkin et al., 2020).

2. **variables extrapolated from digital photography and satellite imagery:**

    (a) tree cover density, derived from a map downloaded from Copernicus Land Monitoring Services, and

    (b) light pollution, extrapolated from night-time digital photographic images shot on 08/10/2015 by astronauts from the International Space Station (Kyba et al., 2015).

Additionally, **spatial variables** including distance to the city centre (Palace of Culture and Science), closest road, and closest path, all measured in metres in QGIS (Corsini et al., 2017).
